# Genetic transformation of the frog-killing chytrid fungus *Batrachochytrium dendrobatidis*

**DOI:** 10.1101/2023.10.17.561934

**Authors:** Erik Kalinka, Stephanie M. Brody, Andrew J. M. Swafford, Edgar M. Medina, Lillian K. Fritz-Laylin

## Abstract

*Batrachochytrium dendrobatidis* (*Bd*), a causative agent of chytridiomycosis, is decimating amphibian populations around the world. *Bd* belongs to the chytrid lineage, a group of early-diverging fungi that are widely used to study fungal evolution. Like all chytrids, *Bd* develops from a motile form into a sessile, growth form, a transition that involves drastic changes in its cytoskeletal architecture. Efforts to study *Bd* cell biology, development, and pathogenicity have been limited by the lack of genetic tools with which to test hypotheses about underlying molecular mechanisms. Here, we report the development of a transient genetic transformation system for *Bd*. We used electroporation to deliver exogenous DNA into *Bd* cells and detected transgene expression for up to three generations under both heterologous and native promoters. We also adapted the transformation protocol for selection using an antibiotic resistance marker. Finally, we used this system to express fluorescent protein fusions and, as a proof of concept, expressed a genetically encoded probe for the actin cytoskeleton. Using live-cell imaging, we visualized the distribution and dynamics of polymerized actin at each stage of the *Bd* life cycle, as well as during key developmental transitions. This transformation system allows, for the first time, direct testing of key hypotheses regarding mechanisms of *Bd* pathogenesis. This technology also paves the way for answering fundamental questions of chytrid cell, developmental, and evolutionary biology.

## INTRODUCTION

Fungal pathogens are a growing threat to human health and global ecology (1). The chytrid fungus *Batrachochytrium dendrobatidis* (*Bd*), for example, has decimated global amphibian populations, endangering the stability of aquatic ecosystems and food webs around the world (2, 3). *Bd* begins life as a motile cell, called a zoospore, that colonizes the skin of frogs, salamanders, and other amphibians (4, 5). The zoospore develops into a multinucleated form called a sporangium that produces new zoospores to spread the infection (4). Although the ecological impact of *Bd* is clear, the molecular mechanisms that drive its developmental program and pathogenesis remain understudied due to a lack of basic molecular genetic tools with which to test core hypotheses about *Bd* biology.

Techniques used to test molecular hypotheses, including CRISPR-mediated genome editing, RNAi-based gene knockdown, and fluorescent protein tagging, are typically based on a robust transformation protocol. Genetic transformation of the soil chytrid *Spizellomyces punctatus* (*Sp*) has recently been used to answer questions about basic chytrid cell biology (6) but, as it is nonpathogenic, cannot be employed to answer questions about infection mechanisms. As the first step toward genetic transformation of *Bd*, Swafford et al. used electroporation to deliver fluorescently-labeled dextrans into cells (7), but did not test whether this protocol could also be used to deliver exogenous DNA for transgene expression in *Bd*.

Here we report the genetic modification of *Bd* by electroporation of plasmid DNA. We show that this transformation is transient, can be used to express a variety of fusion proteins under promoters of various strengths, and provide an example of how the transformation system can be used to study protein distribution during developmental transitions that are key to *Bd* pathogenesis.

## RESULTS

### Electroporation of plasmid DNA yields hygromycin-resistant *Batrachochytrium dendrobatidis* cells

Because zoospores lack cell walls and have only one nucleus (4), we chose this developmental stage for introducing exogenous DNA into *Bd*. To obtain a homogenous population of zoospores, we synchronized *Bd* cultures, resulting in a lifecycle of 3-4 days (**Fig. 1A**) (8). We previously developed an electroporation-based protocol that delivers fluorescently-labeled dextrans to >75% of *Bd* zoospores (7). However, when we attempted to transform *Bd* using this protocol, it consumed an excessive amount of DNA and killed most cells due to higher electric currents caused by the addition of DNA (depressing data not shown). Therefore, we re-optimized this electroporation protocol using smaller cuvettes and a lower-conductivity buffer and achieved high levels of fluorescent dextran delivery into *Bd* zoospores (66%-91% positive cells, **Fig. S1**).

**Figure 1.**
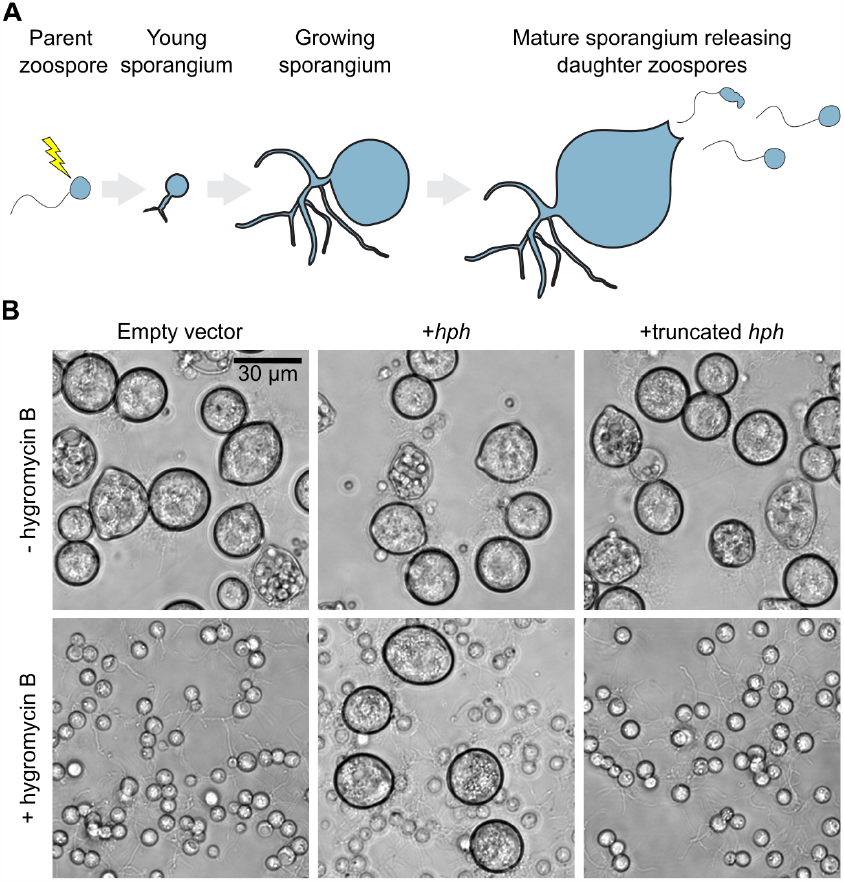
Transient transformation of *Batrachochytrium dendrobatidis* by electroporation of plasmid DNA. (**A**) *Bd* develops from a motile zoospore, then encysts to form a sporangium that then grows for approximately three days before releasing dozens of daughter zoospores. Cell wall-less uninuclear zoospores are used for delivery of plasmid DNA by electroporation. (**B**) Expression of full-length *hph* resistance marker confers transient resistance to hygromycin B. Cells were electroporated with either empty vector (pUC19), full-length *hph* (pEM112B), or truncated *hph* (pEK14) and allowed to recover for one full generation. Daughter zoospores were then transferred into plain media (top row) or media supplemented with hygromycin B (bottom row).

We next used this optimized electroporation protocol to deliver plasmid DNA into *Bd* zoospores. We started with a plasmid that was previously used to successfully transform *Sp* (6). This construct encodes the Hph protein—a selectable marker that confers resistance to hygromycin B, which efficiently kills wild type *Bd* (9)—under the control of the *Sp* histone 2B promoter. We added 40 μg plasmid to 1.2 x 10^6^ zoospores and electroporated the cells. We recovered the cells in non-selective medium for four days until they released daughter zoospores, and then added 0.5 μg/mL hygromycin B to the daughter cells to select for transformants (**Fig. 1B**). Unlike control cells electroporated with an empty vector, which never matured, cells electroporated with the hygromycin resistance marker developed into full-sized sporangia and released zoospores by day 4 of selection (**Fig. 1B**, left and middle columns).

To ensure that the hygromycin resistance was due to the effect of the transgene and not simply a response to stress (10), we also introduced a construct with an early stop codon in the hph gene. Electroporation of this second construct did not confer hygromycin resistance (**Fig 1B**, right column), indicating that resistance conferred by the full length Hph was due to transgene expression. However, all transformants stopped growing by 12 days of selection, hinting that this transformation is transient.

### Transient gene expression persists through at least three lifecycles and can be modulated by use of endogenous *Batrachochytrium dendrobatidis* promoters

To directly test the transience of the transformation, we fused the coding sequence of NanoLuc luciferase, which allows fast and highly sensitive quantification of transgene expression (11, 12), to the C-terminus of the Hph gene. We transformed zoospores on three separate days and measured luciferase activity of zoospore lysates of three successive generations. We observed strong luminescence in the first generation of post-transformation zoospores (**Fig. 2A**). This signal decayed but was still detectable in the second and third generation when compared to the empty vector controls (for example, 15.8 +/- 9.7 fold on day 12). Quadrupling the amount of DNA led to a reproducible increase in luminescence (3.2 +/- 1.0 fold on day 4) but did not lead to stable luciferase activity.

**Figure 2.**
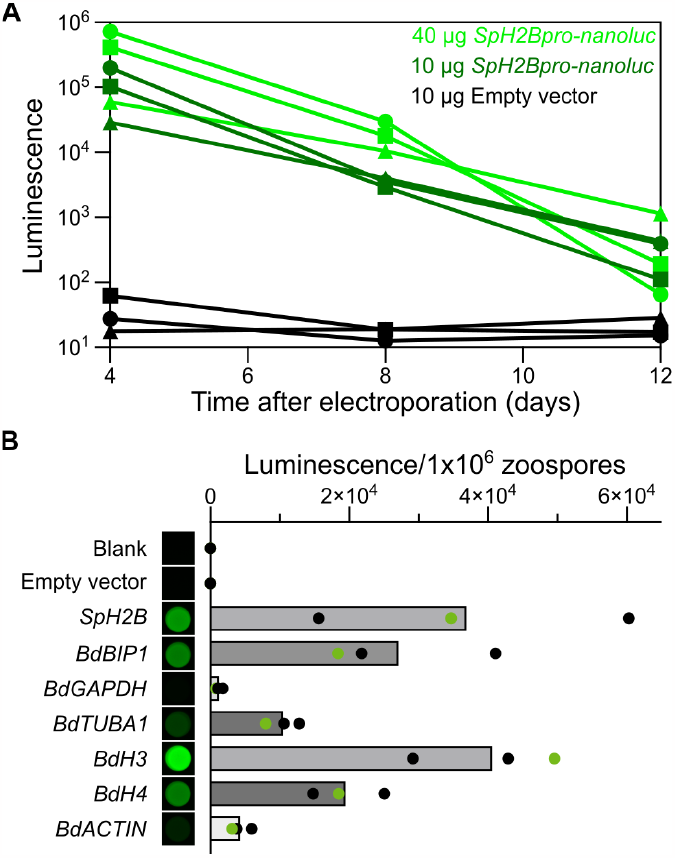
Luciferase reporter assay facilitates quantification of heterologous gene expression in *Batrachochytrium dendrobatidis*. (**A**) Luciferase expression is transient and signal levels depend on the amount of transfected reporter plasmid. Signal intensity was normalized to the number of transfected cells used for each timepoint. Data from three independent trials is shown with individual trials indicated by different symbols. (**B**) NanoLuc fusions were used to quantify transgene expression from various promoters. *Bd* zoospores were electroporated with 40 μg of each reporter plasmid and assayed for luciferase expression after 4 days of recovery. NanoLuc luminescence were detected in well plates using a gel imager (left) and quantified using a plate reader (right). Circles represent independent replicates and grey bars the mean. Green circles indicate plate reader quantification of samples shown on the left. Plasmids used: empty vector (pUC19), *SpH2B* (pGI3EM17), *BdBIP1* (pEK19), *BdGAPDH* (pEK20), *BdTUBA1* (pEK21), *BdH3* (pEK22), *BdH4*(pEK23), *BdACTIN* (pEK24).

Although the *Sp* H2B promoter clearly allows transgene expression in *Bd*, to fine-tune expression levels, we tested different endogenous promoters to drive expression of the Hph-NanoLuc fusion. We selected *Bd* homologs of genes whose promoters are commonly used to drive expression in established model systems and inserted up to 1 kb of their 5’ noncoding sequence upstream of the Hph-NanoLuc fusion (**Table S1**). Following electroporation, all sequences drove gene expression at levels detectable by luciferase assays and conferred transient resistance to hygromycin B (**Fig. 2B**). The luminescence intensity of each construct was at least two orders of magnitude greater than that of cells transformed with an empty vector (**Fig. 2B**); while the GAPDH, actin, and α-tubulin promoters yielded relatively low levels of expression, the promoters of histone 4, histone 3, and BiP1 showed high levels of expression.

### Detection of co-transformed fluorescent proteins in *Batrachochytrium dendrobatidis* cells

We next adapted the transformation protocol for expressing fluorescent protein fusions. To be detected, fluorescent proteins must be expressed at much higher levels than enzymes like NanoLuc. Because the *Sp* H2B promoter gave consistently high expression (**Fig. 2**), we first tested the plasmids previously established for fluorescent protein expression in *Sp* (6). We transformed constructs encoding mCerulean3, mClover3, and tdTomato into *Bd* zoospores and observed cytosolic fluorescence of the expected wavelengths (**Fig. 3A, Fig. S2A**). We also engineered a new construct encoding a monomeric red fluorescent protein, mRuby3 (13), that gave similar results (**Fig. 3A**).

**Figure 3.**
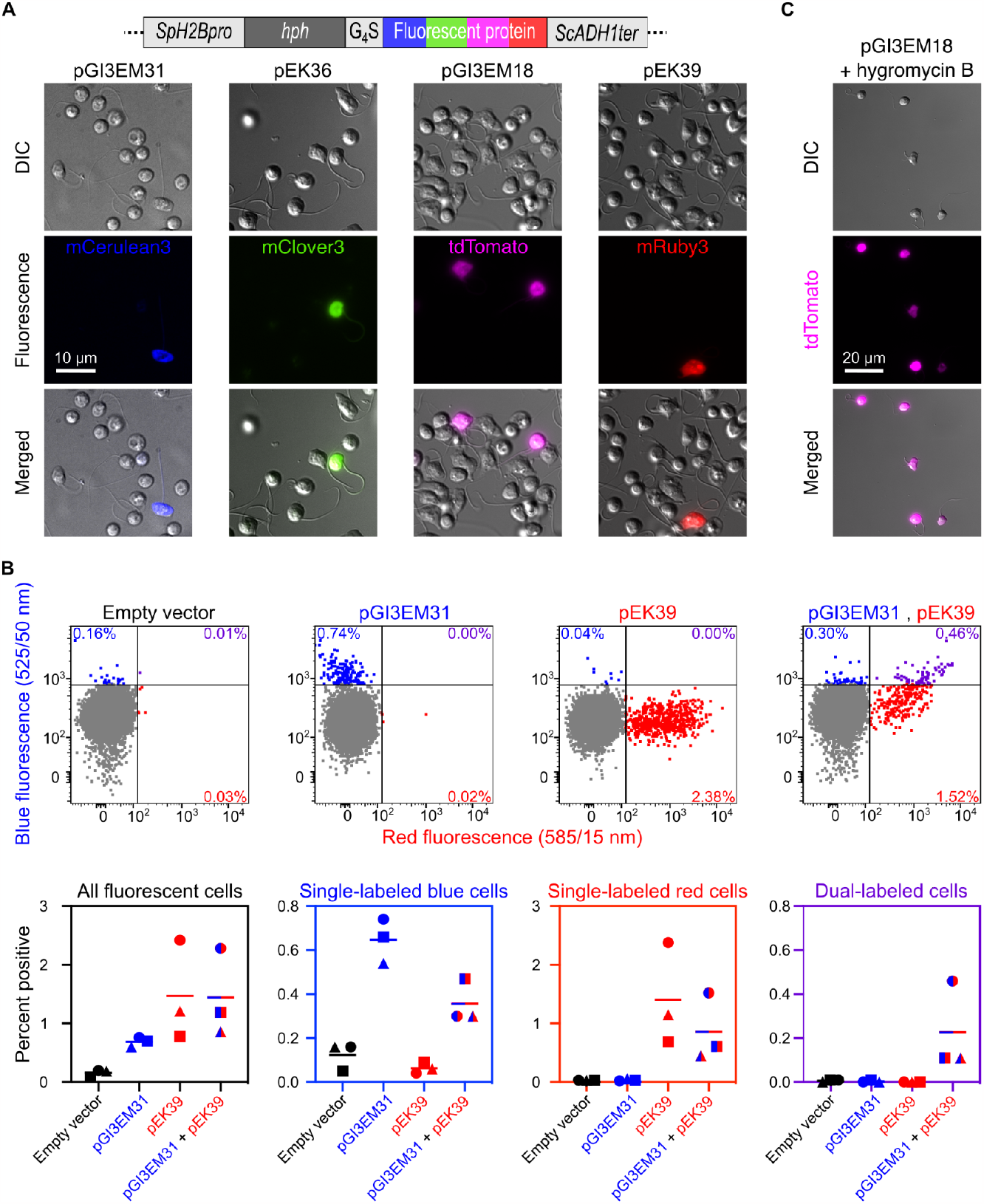
Diverse fluorescent proteins are functional in *Batrachochytrium dendrobatidis*. (**A**) Expression of different fluorescent fusion proteins in *Bd* zoospores. *Bd* zoospores were electroporated with 40 μg of each indicated plasmid, and their daughters were imaged using widefield and DIC microscopy. (**B**) Transformation efficiency was measured by flow cytometry. *Bd* zoospores were electroporated with empty vector (pUC19), or plasmids encoding mCerulean3 (pGI3EM31) and/or mRuby3 (pEK39), and their daughter cells then analyzed using flow cytometry. The top row shows red and blue fluorescence intensities of cell populations from one of three independent experimental replicates. Bottom row shows quantification of the percent of fluorescence cells from all three replicates. Replicates are represented by different symbols and bars indicate means. (**C**) Cells were enriched for fluorescent protein expression by early addition of hygromycin B to the medium. Hygromycin B was added to cells transformed with the tdTomato plasmid (pGI3EM18) 18 hours after electroporation and the next generation of zoospores imaged as in (A). Control cells electroporated with the tdTomato plasmid that were grown without hygromycin B are shown in (A).

We next quantified the transformation efficiency and tested whether we could successfully co-transform multiple constructs. We transformed *Bd* zoospores with mCerulean3 and/or mRuby3 encoding constructs and used flow cytometry to measure the percent of cells with detectable blue and red fluorescence (**Fig. 3B**). We found that transforming cells with a single construct gave ∼1% positive cells (0.6 +/- 0.1% blue positive cells for mCerulean3 and 1.4 +/- 0.9% red positive cells for mRuby3, **Fig. 3B**, bottom column). Transforming with both constructs yielded 0.1-0.5 % dual-fluorescent cells, indicating that ∼15% of cells that receive the mRuby3 construct also receive the mCerulean3 construct (**Fig. 3B**). Because only a small percentage of the cell population in single and/or double transformations showed detectable fluorescence, localization studies using this approach would be tedious. We therefore tested whether early application of selection pressure could enrich fluorescent cells. Adding hygromycin B to cells 18 hours post electroporation resulted in the majority of cells displaying detectable fluorescence (**Fig. 3C**). Based on these data, we recommend application of selection 18 hours after electroporation for quick identification of transformants.

Fluorescent protein expression often increases upon codon optimization (14). Thus far, all of the transgenes we tested in *Bd* were codon-optimized for mammalian cell expression because this approach was successful for *Sp*. However, we recognized that the frequency of codons in the *Bd* genome is not similar to that of mammals, but is close to that of *Saccharomyces cerevisiae* (**Data Set 1**). Because yeast codon-optimized sequences are readily available, we therefore tested whether yeast codon usage could increase expression levels in *Bd* by engineering matched constructs encoding Hph-mClover3 and Hph-mRuby3 adapted for *Homo sapiens* or *S. cerevisiae* codon usage (**Fig. S2B**-**C**). Surprisingly, only the constructs codon-optimized for *H. sapiens* were functional in *Bd* (**Fig. S2D**). Although the mechanisms underlying this phenomenon remain unclear, we continued to use constructs optimized for mammalian cell expression for *Bd* transformation.

### Transient transformation of *Batrachochytrium dendrobatidis* reveals actin dynamics during key developmental transitions

Finally, to illustrate how transient transformation can be used to study *Bd* development and pathogenesis, we expressed and imaged a fluorescent cytoskeletal marker. In many eukaryotic organisms, including other chytrid species (6), dynamic remodeling of the actin cytoskeleton drives developmental transitions (15). We previously used fixed cells to show that *Bd’s* actin cytoskeleton is highly variable (16–18). For example, zoospores build a layer of actin that supports the cell membrane from beneath called an actin cortex. Zoospores also assemble broad actin-filled protrusions called pseudopods that we inferred to be involved in cell crawling (16), a hypothesis that we could not directly test without imaging actin dynamics in living cells. In contrast, sporangia form actin shells around their nuclei, as well as actin patches at the cell periphery (17). We therefore hypothesize that the developmental transition from zoospore to sporangium, called encystation, involves drastic rearrangement of the actin cytoskeleton. To directly test these two hypotheses we transformed *Bd* with tdTomato fused to LifeAct, a small actin-binding peptide, and visualized cytoskeletal dynamics during zoospore crawling and encystation (6, 19).

To confirm the fidelity of LifeAct localization in *Bd*, we fixed and stained wild type and transformed cells with fluorescent phalloidin, a small molecule that binds to actin polymer with high specificity (20). In contrast to the cytosolic localization of tdTomato alone, LifeAct-tdTomato co-localized with phalloidin in both zoospores (**Fig. 4A**) and sporangia (**Fig. 4B**). In zoospores, both phalloidin and LifeAct revealed broad actin-filled pseudopods and a thick actin cortex (**Fig. 4A**), a co-localization that we confirmed using line scans (**Fig. 4A, bottom**). In sporangia, phalloidin and LifeAct-tdTomato co-localized to actin shells around the nuclei and actin patches at the cell periphery (**Fig. 4B**), consistent with previous observations (17). These data indicate LifeAct-tdTomato specifically highlights actin structures throughout the *Bd* lifecycle.

**Figure 4.**
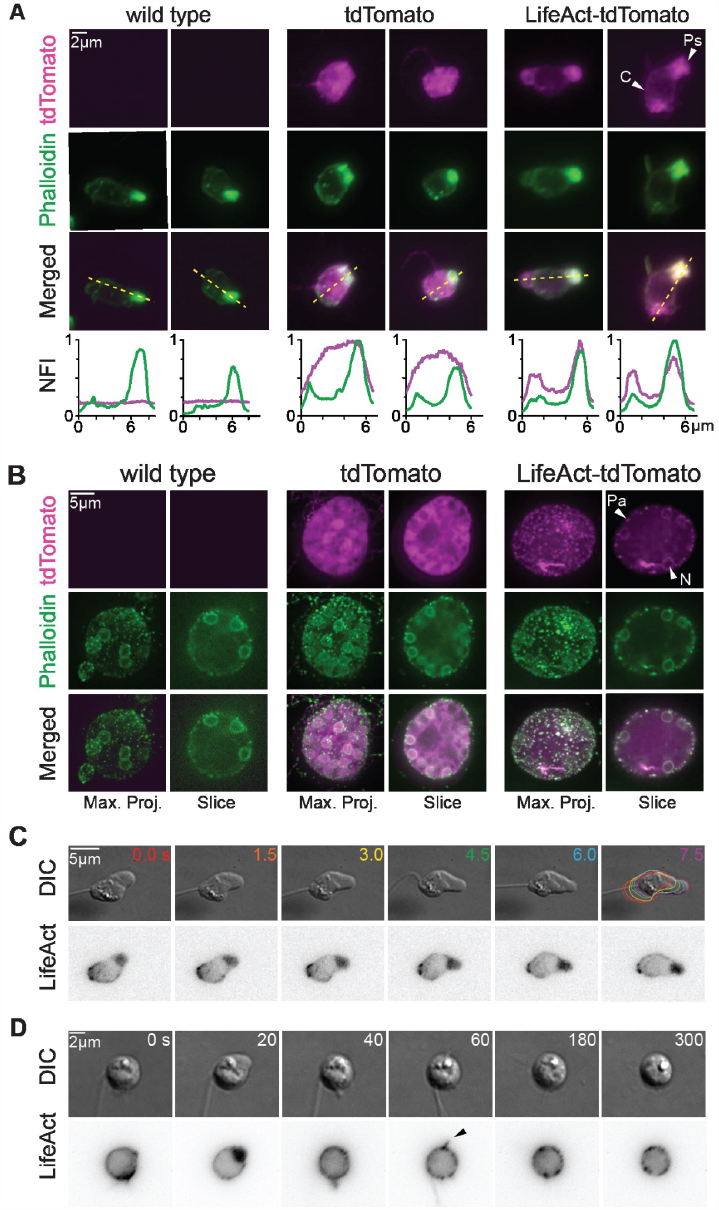
Localization of LifeAct-tdTomato visualizes the distribution and dynamics of polymerized actin in live *Batrachochytrium dendrobatidis* cells. (**A**) Representative maximum intensity projections of two wild type (left), two tdTomato (center, pGI3EM18) and two LifeAct-tdTomato (right, pGI3EM22C) expressing zoospores fixed and stained with phalloidin (green). Graphs show normalized fluorescence intensity (NFI) of phalloidin (green) and 561 nm (magenta) along shown dotted yellow lines. White arrows point to examples of a pseudopod (Ps) and an actin cortex (C). (**B**) Representative maximum intensity projections and single slice images of wild type (left), tdTomato (center) and LifeAct-tdTomato (right) expressing sporangia fixed and stained with phalloidin (green). White arrows point to examples of an actin patch (Pa) and a nuclear shell (N). (**C**) Zoospore expressing LifeAct-tdTomato crawling in confinement. Colored cell outlines highlight movement over time. (**D**) Zoospore expressing LifeAct-tdTomato undergoing actin cytoskeletal remodeling upon mucin treatment. Image sequence starts 3 minutes after mucin treatment. Black arrow points to a thin, actin-filled protrusion.

We next used LifeAct-tdTomato to visualize actin dynamics in living *Bd* cells, starting with crawling zoospores. Time-lapse imaging of these cells revealed dynamic pseudopods at the leading edge that were filled with dense networks of actin (**Fig. 4C, Video S1**), confirming our previous hypothesis of how *Bd* zoospores crawl (16). To test the hypothesis that *Bd* development involves rapid actin cytoskeleton remodeling, we imaged cells during encystation. We recently showed that *Bd*, which grows within the mucus-coated skin of amphibians, rapidly encysts upon exposure to mucin, the primary component of mucus (18). To visualize changes in the actin cytoskeleton during encystation, we therefore imaged LifeAct-tdTomato-expressing zoospores while exposing them to mucin. Within several minutes of mucin treatment, zoospores stopped forming pseudopods, disassembled their actin cortex, and formed actin patches at the cell periphery (**Fig. 4D, Video S2**). This rapid transition from zoospore-associated actin structures to sporangium-associated actin structures confirms our hypothesis that encystation involves rapid actin remodeling. During this transition, we also observed thin protrusions that contained LifeAct. These actin structures were not previously reported, perhaps because they are transient and inconspicuous when imaged with DIC alone (**Fig. 4D, Video S2**). Taken together, these examples demonstrate the use of fluorescent protein expression to test hypotheses about the development and pathogenesis of *Bd*.

## DISCUSSION

By developing a transient transformation protocol for *Bd*, we have established this frog-killing fungus as a genetic model system for chytrid biology and pathogenesis. To validate its efficacy, we quantified transgene expression through multiple *Bd* generations under the control of various endogenous promoters. We detected expression through three generations, but not beyond that point. We suspect that this transience is due to plasmids being diluted across generations rather than being replicated episomally or being integrated into the host genome. This effect may be accelerated compared to other organisms because each chytrid life cycle involves production of dozens of daughter cells from a single parent (21). Because *Bd* does not appear to readily integrate heterologous DNA into its chromosomes, stable transformation may require designing vectors that can exploit native homologous recombination pathways.

This transient transformation system opens the door to testing molecular hypotheses about *Bd* pathogenesis. By expressing fluorescent protein fusions, for example, we can explore the dynamics of endogenous *Bd* proteins during development and infection. We previously had indirect evidence that *Bd*’s actin cytoskeleton undergoes drastic remodeling throughout its lifecycle (16–18). Here we confirmed this hypothesis by transforming cells with a fluorescently labeled actin probe to directly visualize changes in the actin cytoskeleton during key developmental transitions. This tagging approach can also be used to study proteins that have been hypothesized to play specific roles in *Bd* pathogenesis, such as metalloproteases (22), chitin binding proteins (23), and Crinkler effector proteins (24). To directly test the function of these or any *Bd* protein, however, we need techniques for modulating endogenous gene expression. To this end, transient transformation is well-suited for CRISPR-mediated genome editing because continuous expression of Cas enzymes can be cytotoxic (25). Alternatively, as chytrids have the genes required for RNAi (26), transformation could be used to express dsRNA to initiate RNAi-mediated gene silencing. These and other genetics tools lay the foundation for understanding *Bd* pathogenesis and for developing strategies to mitigate the effects of chytrid disease.

This transient transformation system also relieves a major bottleneck to evolutionary studies. Due to their early divergence, chytrids are widely used to study deep fungal evolution (21). There are over a thousand chytrid species, of which *Bd* is only the second to become genetically tractable after the nonpathogenic species *Spizellomyces punctatus* (6). *Sp* and *Bd* diverged from a common ancestor ∼500 million years ago (27), and have since evolved differences in their basic biology (17, 21), as well as in pathogenesis. With the advent of genetic engineering in *Bd*, along with the recent transformation of *Sp*, we can now begin to investigate the molecular mechanisms that shaped early fungal evolution.

The genetic diversity of chytrids is on par with that of animals (28). These species inhabit diverse ecosystems and take part in a wide variety of symbioses. To understand how chytrids have adapted to these novel lifestyles, we need additional genetically tractable species. Because electroporation-based DNA delivery is readily adaptable to other species (7), we look forward to tailoring these tools to other ecologically-relevant chytrid fungi.

## MATERIALS AND METHODS

### Cell culture

We used the *Batrachochytrium dendrobatidis* strain JEL423 (from the highly virulent global panzootic lineage) for all experiments. Unless stated otherwise, *Bd* cells were grown in 1% tryptone (w/v) at 24 °C in tissue culture-treated flasks (Fisher, Cat. No. 50-202-081). Cultures were synchronized and split every three days by collecting released daughter zoospores and using them to seed the next generation in a fresh flask. For additional details, see (8).

### Reporter plasmid design and molecular cloning

We generated new expression constructs by SLIC (29, 30) and seamless cloning (NEBuilder® HiFi DNA Assembly, New England Biolabs, Cat. No. E2621S). For PCR amplification from plasmids or *Bd* genomic DNA, we used the Phusion high-fidelity DNA polymerase (New England Biolabs, Cat. No. M0530L). We sequence-verified all constructs by whole plasmid sequencing (SNPsaurus LLC). *Saccharomyces cerevisiae* and *Homo sapiens* codon-optimized reporter genes were synthesized by Twist Bioscience (South San Francisco, United States). For transformation experiments, we performed large scale preparations of reporter plasmids with a Midiprep kit (NucleoBond Xtra Midi, Machery-Nagel, Cat. No. 740410.50) and resupsended the DNA pellet in low TE buffer (10 mM Tris, 0.1 mM EDTA, pH 8.0) to a concentration of at least 2.7 μg/μL. For information about each plasmid used in this study, see **Data Set 2**. For testing each native *Bd* promoter, with the exception of *Bd* actin, we cloned the entire 5’ intergenic region, including the putative promoter and 5’UTR, from the target genes into *nanoluc* reporter plasmids (**Table S1**). For the *Bd* actin promoter, we cloned 1010 bp immediately upstream of the translation start site. Complete plasmid sequences and plasmids have also been deposited at Addgene (https://www.addgene.org/Lillian_Fritz-Laylin/, **Data Set 2**).

### Optimized electroporation

We optimized the following electroporation protocol for 1 mm electroporation cuvettes and delivery of plasmid DNA into *Bd* zoospores. We sterilized all solutions and conducted all steps of the transformation protocol in a biosafety cabinet. *Bd* zoospores were synchronized as previously described (7). When working with a healthy *Bd* cell culture, a 175-cm^2^ flask yields enough synchronized zoospores for 3-4 electroporation cuvettes.

On the day of electroporation, we prepared fresh electroporation buffer AS27 (25 mM D-Mannitol, 0.7 mM MgCl_2_, and 1 mM KCl) and sample working solutions (fluorescent dextrans or reporter DNA). 50 mg/mL stock solutions of fluorescein-labeled dextrans (Invitrogen, Cat. No. D3305), were prepared in ultrapure water and diluted in AS27 to a final concentration of 2 mg/mL. Reporter plasmids were first diluted into 15 μL of low TE buffer, and then mixed with 25 μL of AS27 in a microcentrifuge tube. Additional AS27 buffer, sample working solutions, 1% tryptone media, and 1 mm electroporation cuvettes (Bulldog Bio, Cat. No. 12358-345) were chilled on ice.

We collected synchronized *Bd* zoospores in media and strained them through a sterilized 25 mm Whatman Grade 1 filter paper (CAT No. 1001–325) to remove sporangia and cell clumps. We then washed the zoospores three times with 10 mL of chilled AS27 by centrifugation at 2,500×g for 5 minutes at 4°C, then resuspended the cells in 100 μL of AS27 and measured zoospore density using a hemocytometer and brought the zoospore suspension to a working concentration of 3x10^7^ cells/mL with AS27 buffer.

To electroporate the zoospores, we added 40 μL of zoospore cell suspension and 40 μL of sample working solution to the prechilled cuvettes without mixing. We then placed the cuvettes on ice for 10 minutes before electroporating. Just before inserting each cuvette into the electroporator, we mixed the contents by pipetting up and down 3-4 times. We then placed the cuvette in the ‘shock pod’ of the electroporator (Nepa Gene ELEPO21), measured the impedance, and electroporated the cells using two positive polarity 350 V/0.2 ms poring pulses at 50 ms intervals, followed by one 20 V/50 ms polarity-exchanged transfer pulse at 50 ms intervals. The impedance of a transformation mixture containing 40 μg DNA should be 400 to 600 Ω, giving off 0.08 to 0.11 J energy by the poring pulses. Immediately after pulsation, we placed the cuvettes on ice and allowed them to rest for 10 min, after which we added 200 μL of ice-cold 1% tryptone without mixing. Following another 10 min of incubation on ice, we added an additional 400 μL of ice-cold tryptone to the cuvette, mixed the cell solution by pipetting up and down several times, and transferred the cells to a 5 mL microcentrifuge tube that contained 3.9 mL chilled tryptone. When working with dextran samples, we prepared the cells for flow cytometry at this point. For cells electroporated with plasmids, we split each sample into 600 μL aliquots and transferred these to tissue culture-treated 24-well plates (Celltreat, Cat. No. 229124).

### Antibiotic resistance

When early application of selection pressure was desired (**Fig. 3C, Fig. 4**), we allowed the cells to recover for 18 h at 24°C, and then added hygromycin B (Gibco, Cat. No. 10687010) dissolved in 1% tryptone to each well to give a final concentration of 0.5 μg/mL, and incubated the cells at 24°C for 78 h. Otherwise (**Fig. 1B**), we grew the cells at 24°C for 4 days before assaying them for hygromycin B resistance by pooling the cells from each sample, filtered them through a sterilized 25 mm Whatman Grade 1 filter paper, centrifuged them for 5 min at 2,500×g at room temperature, and finally resuspended them in fresh 1% tryptone supplemented with 0.5 μg/ml hygromycin B. We then incubated the cells for 4 days at 24°C and analyzed their growth using light microscopy.

### Nanoluc reporter assay

We collected zoospores released from electroporated cell populations, filtered them through a 25 mm Whatman Grade 1 filter paper, and removed 10 μL to count the number of cells for normalization. We then pelleted the rest of the sample by centrifugation for 5 min at 2,500xg at room temperature and resuspended the cell pellet in 50 μL of Bonner’s salts (10.27 mM NaCl, 10.06 mM KCl, 2.7 mM CaCl_2_ in ultrapure water). We then mixed the cell solution with 80 μL of NanoGlo luciferase assay reagent (Promega, Cat. No. N1110), transferred it to a 96-well plate well (PerkinElmer,Cat. No. 6005290), and measured luminescence using a FilterMax F5 microplate reader (Molecular Devices) every minute for 1 h using a 400 ms integration time. Reported luminescence values represent the mean of the measurements after 9, 10, and 11 minutes. Immediately after the plate reader measurement, we used a G:BOX iChemi XR gel imaging system (Syngene) to visualize NanoLuc luminescence in individual wells.

### Flow cytometry

For dextran electroporation, we washed cells thrice with Bonner’s salts by centrifugation at 2,500xg for 5 min at room temperature, and resuspended the zoospore pellet in 600 μL fresh Bonner’s salts, transferred them to a fresh microcentrifuge tube and added 300 μL paraformaldehyde (PFA) based fixation buffer to give a final concentration of 4% PFA, 45 mM sucrose, 25 mM sodium phosphate buffer (pH 7.2). We incubated the zoospores for 20 min on ice, and washed them with 400 μL unchilled Bonner’s salts. For plasmid electroporation, we collected the first post-electroporation generation of zoospores, filtered them through 25 mM Whatman Grade 1 paper, and washed and fixed the zoospores similar to dextran-treated cells, but used a diluted fixation buffer to reduce quenching of protein fluorescence (final concentration: 1.6% PFA, 30 mM sucrose, 16 mM sodium phosphate buffer, pH 7.2). We performed all flow cytometry measurements on a BD Dual LSRFortessa cell analyzer (BD Biosciences) using the 405, 488, and 561 nm lasers to detect mCerulean3, FITC, and mRuby3 fluorescence, respectively. We captured 10,000 events for dextran measurements and a minimum of 30,000 events for fluorescent protein detection in transformants. Compensation of multi-color experiments was conducted using the BD FACSDiva software (BD Biosciences). We analyzed the resulting data using FlowJo v10.9, identifying singlets using the forward scatter area versus height signals and setting gates based on data recorded from cells electroporated without a fluorescent reporter.

### Cell fixation and staining

Zoospores were adhered for 5 min to plasma cleaned 96-well plates (MatriPlate, Cat. No. MGB096-1-2-LG-L) coated with 0.5 mg/mL Concanavalin A (Sigma, Cat. No. C2010). We fixed the cells in 4% PFA and 50 mM sodium cacodylate, pH 7.2 on ice for 20 min, then washed them with PEM (100 mM PIPES, pH 6.9, 1 mM EGTA, 0.1 mM MgSO_4_). We then permeabilized and stained the cells with 66 nM Alexa Fluor 488 Phalloidin (Thermo Fisher, Cat. No. A12379) in 0.1% Triton X-100 PEM buffer for 20 min at room temperature, and washed once with PEM.

### Cell crawling assay

To confine cells, we used a one-well dynamic cell confiner with a suction cup and 2 μm confinement slides from 4Dcell (Montreuil, France). To hydrate the device, we incubated the suction cup and confinement slides in Bonner’s salts for at least 1 h before use. We seeded 150 μL of zoospores suspended in Bonner’s salts into a single well of a 6-well glass bottom plate (Fisher Scientific, Cat. No. NC0452316) and allowed the cells to settle for 10 min before gradually applying confinement by decreasing the pressure from -3 kPa to -10 kPa.

### Mucin-induced encystation

Zoospores were adhered to plasma cleaned 96-well plates coated with 0.5 mg/mL Concanavalin A. We resuspended mucin (Sigma, Cat. No. M1778) in Bonner’s salts at 20 mg/mL and centrifuged the solution at 15,000×g for 5 minutes to remove particulates and treated adhered zoospores with mucin at a final concentration of 10 mg/mL.

### Microscopy

For experiments shown in **Fig. 1B** and **Fig. S2D**, we used a Nikon Ti2-E inverted microscope, equipped with a 40x Plan Fluor 0.6 NA objective (with a correction collar set to 1.2) and a sCMOS 4mp camera (PCO Panda) controlled through NIS-Elements software (Nikon). We acquired images using both Phase-contrast (white LED transmitted light) and epifluorescence microscopy with LED illuminators at 460 nm to visualize mClover3, and 550 nm to visualize mRuby3. We used the same microscope with a Plan Apo λ 100x 1.45 NA oil objective for all cells shown in **Fig. 3, Fig. 4A, Fig. 4C-D**, **Video S1, Video S2**, and **Fig. S2A** except for mCerulean3-expressing cells displayed in **Fig. 3A**. We took images using differential interference contrast (DIC, acquired using transmitted light) and epifluorescence microscopy with LED excitation light at 460 nm to visualize mClover3 and Alexa Fluor 488 phalloidin, and 550 nm to visualize mRuby3 and tdTomato. For z-stack imaging of entire zoospores, we acquired single plane images with 0.2 μm steps. For images of sporangia shown in **Fig. 4B** and mCerulean3-expressing zoospores shown in **Fig. 3A**, we used a Nikon Ti2-E microscope, equipped with a Plan Apo λ 100x 1.45 NA oil objective, a Crest X-Light spinning disk (50 μm pinhole), a Photometrics Prime 95B camera, a Lumencor Celesta light source for fluorescence, controlled through NIS-Elements AR software. We acquired images using DIC and fluorescence microscopy with excitation light via a 446 nm laser to visualize mCerulean3, 477 nm laser to visualize Alexa Fluor 488 phalloidin, and 546 nm laser to visualize tdTomato. For z-stack imaging of entire sporangia, we acquired single plane images with 1 μm steps. We performed all imaging at room temperature, with live cells suspended in Bonner’s salts and fixed cells in PEM buffer, and all image analysis in Fiji (31).

## Supporting information

Data Set 1

Data Set 2

## ACKNOWLEDGMENTS

We thank Tim James and Fritz-Laylin lab members for comments on the manuscript and David Booth for his fantastic manuscript on *Choanoflagellate* transformation that convinced us to try using NanoLuc. This work was funded by the Gordon and Betty Moore Foundation (award #9337), the National Science Foundation (IOS-1827257), and a Pew Scholar award from the Pew Charitable Trusts to L.K.F.-L., who is a CIFAR fellow in the Fungal Kingdom: Threats and Opportunities program.

## SUPPORTING INFORMATION

**Table S1.** Promoters used in *nanoluc* reporter plasmids. For *Bd* promoters, non-coding sequences were amplified from *Bd* JEL423 genomic DNA and cloned upstream of hph-nanoluc fusions. For the *Sp* H2B promoter, the sequence identified by Medina et al. (6) was used.

| Name | Gene description | *nanoluc* reporter plasmid | NCBI Accession Number | Length of cloned 5' noncoding region (bp) |
| --- | --- | --- | --- | --- |
| *SpH2B* | *Sp histone H2B* | pGI3EM17 | XM_016750634.1 | 283 |
| *BdBIP1* | *Bd binding immunoglobulin protein 1, heat shock protein 70 family protein* | pEK19 | XM_006675097.1 | 365 |
| *BdGAPDH* | *Bd glyceraldehyde-3-phosphate dehydrogenase* | pEK20 | XM_006681561.1 | 271 |
| *BdTUBA1* | *Bd alpha-tubulin* | pEK21 | XP_006675832.1 | 930 |
| *BdH3* | *Bd histone H3* | pEK22 | XP_006680289.1 | 298 |
| *BdH4* | *Bd histone H4* | pEK23 | XP_006680247.1 | 298 |
| *BdACTIN* | *Bd actin* | pEK24 | XP_006676494.1 | 1010 |

**Figure S1.**
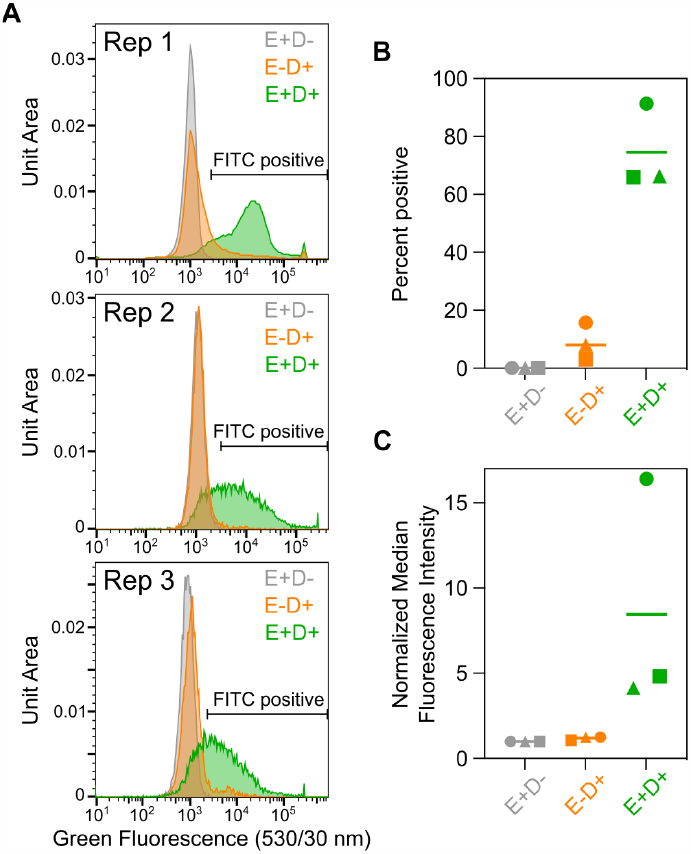
Efficient electroporation and loading of *Batrachochytrium dendrobatidis zoospores* with FITC-dextran. (**A**) Flow cytometry data of the reoptimized electroporation protocol using the ELEPO21 electroporator and AS27 buffer from three independent biological replicates. Shown are: green fluorescence intensity of single cell events for electroporated, no dextran controls (E+D-, grey); non-electroporated, FITC-dextran incubated controls (E-D+, orange); and electroporated, FIC-dextran incubated samples (E+D+, green). Gates for FITC positive cells were set using the E+D-controls for each replicate. (**B**) Percent of FITC positive cells from replicates shown in (A). (**C**) Median green fluorescence intensity of single cell events shown in (A), with values normalized to the median intensities of the electroporated, no dextran controls (E+D-) which are set to 1 for each replicate. Replicates are represented by different symbols and bars indicate means.

**Figure S2.**
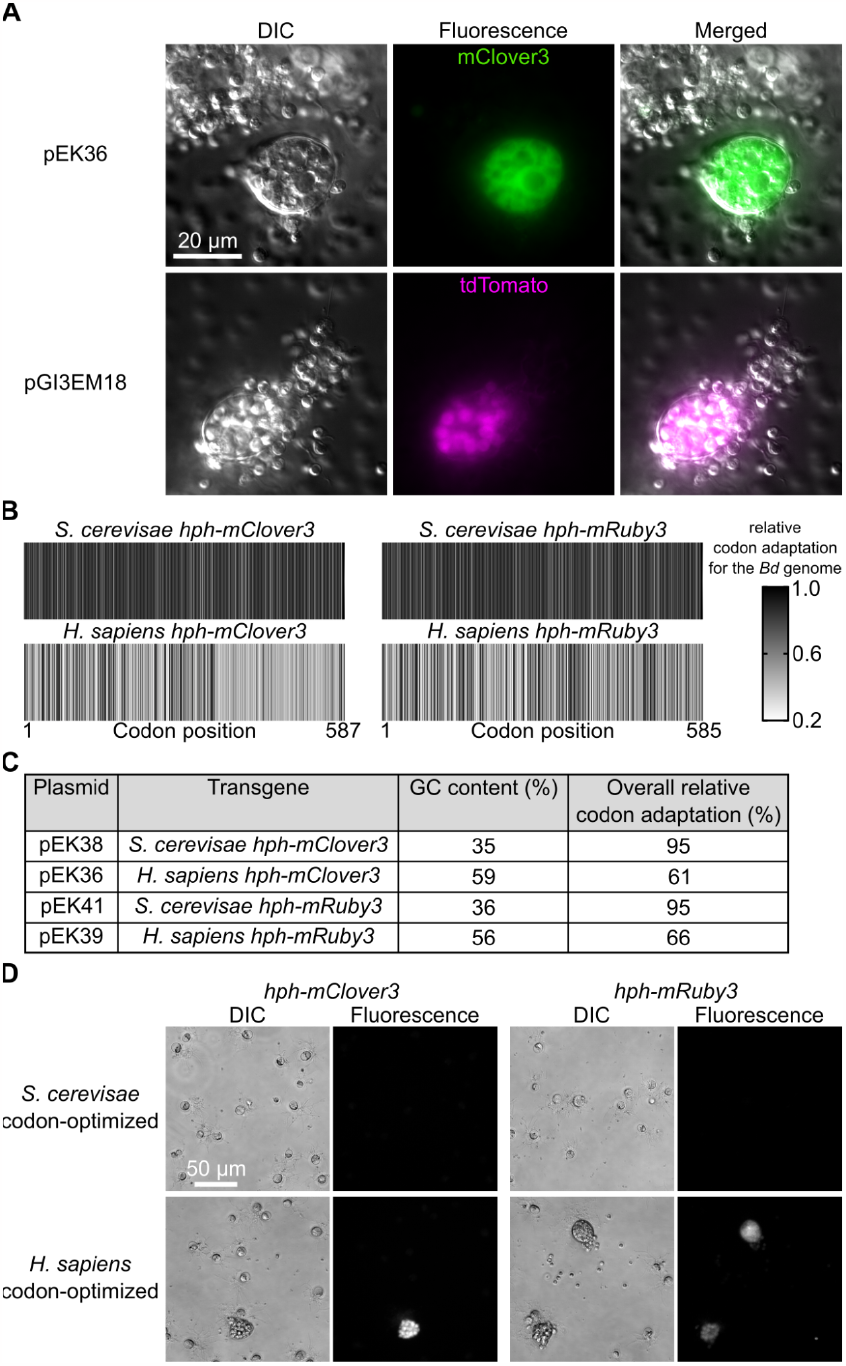
Influence of codon usage on the expression of transgenes in *Batrachochytrium dendrobatidis*. (**A**) Expression of fluorescent proteins in *Bd* sporangia. Zoospores were electroporated with 40 μg of each indicated plasmid, allowed to grow for four days, then imaged using widefield and DIC microscopy. (**B**) Relative codon adaptation of the different fluorescent marker gene variants compared with the nuclear genome of *Batrachochytrium dendrobatidis*. Coloring of bars indicates the relative adaptation of each individual codon across the reading frame of each sequence. (**C**) Key properties of the different fluorescent marker gene variants. Variants of the same gene have the same size and identical amino acid sequence, but vary in their GC content and codon usage. (**D**) Epifluorescence microscopy of cells electroporated with plasmids described in (B) and (C). *Bd* zoospores were transformed with 40 μg of each indicated construct. 18 hours later, 0.5 μg/mL hygromycin B was added and cells were imaged four days after electroporation.

## SUPPLEMENTARY VIDEO AND DATA SET LEGENDS

**Video S1. Video of LifeAct-tdTomato-expressing zoospore in Figure 4C crawling**. Images were taken every 1.5 seconds in DIC (left) and tdTomato (right). Time is displayed in minutes:seconds.

**Video S2. Video of LifeAct-tdTomato-expressing zoospore in Figure 4D undergoing mucin-induced encystation**. Video starts 3 minutes after mucin treatment. Images were taken every 5 seconds in DIC (left) and tdTomato (right). Time is displayed in minutes:seconds.

**Data Set 1**. Codon Usage Comparison between *Batrachochytrium dendrobatidis* (*Bd*), *Saccharomyces cerevisiae* (*Sc*), and *Homo sapiens* (*Hs*). This data set shows the relative adaptiveness (32) for each codon and the corresponding percent deviation (i.e. the difference between the relative adaptiveness of the two indicated species multiplied by 100). Deviation values are highlighted with a color scale, where blue indicates low deviation, white indicates the median, and red indicates high deviation. Codon usage tables were sourced from the HIVE-CUTs databases (33).

**Data Set 2. *Batrachochytrium dendrobatidis* expression plasmids used in this study**. This data set lists the names, source, and cloning strategy used for each plasmid.

## Notes

### Competing Interest Statement

The authors have declared no competing interest.

